# Evaluation of Morpho-physiological Traits of F_1_ Hybrids and Estimation of Standard Heterosis of Sweet Pepper (*Capsicum annuum* L.) in Sylhet

**DOI:** 10.1101/2025.09.18.677104

**Authors:** Abdullah Al Kafi, Jannatul Ferdousi, Md. Iqbal Hossain, Md. Shahidul Islam, Dwipok Deb Nath, Md. Abdul Karim, Swapan Kumar Roy

**Author notes:** Corresponding Author: Dr. Jannatul Ferdousi. **Co-Corresponding author:** Dr. Swapan Kumar Roy. 1 Abdullah Al Kafi. 2 Jannatul Ferdousi. 3 Md. Iqbal Hossain. 4 Md. Shahidul Islam. 5 Dwipok Deb Nath. 6 Md. Abdul Karim. 7 Swapan Kumar Roy.

## Abstract

Sweet pepper (*Capsicum annuum* L.) is a highly demanded crop valued for its culinary and nutritional benefits. This study aimed to evaluate the morphological, physiological, and yield-contributing traits of developed hybrids and check varieties, as well as estimate standard heterosis. The experiment was conducted at the Horticulture Department, Sylhet Agricultural University, from October 2022 to June 2023, using eight hybrids (P_3_×P_1_, P_4_×P_3_, P_6_×P_5_, P_5_×P_1_, P_7_×P_1_, P_5_×P_3_, P_3_×P_2_, and P_6_×P_3_) and two check varieties (CV_1_ and CV_2_). The field and laboratory experiments followed RCBD and CRD designs, respectively, with three replications. Significant morphological variations were observed in growth habit, leaf shape, flower position, fruit color, size, and nodal anthocyanin. The hybrids P_5_×P_1_, P_7_×P_1_, P_6_×P_5_, P_3_×P_2_, and P_5_×P_3_ exhibited superior morphological, physiological, and yield traits compared to check varieties. Earliness in flowering (30.33 days) and fruit setting (38.33 days) was recorded in P_3_×P_1_. Maximum fruit length (13.57 cm) was observed in P_5_×P_1_, while the highest fruit diameter (8.71 cm) and individual fruit weight (214.51 g) were found in P_3_×P_2_. The highest fruit yield per plant (1.94 kg) and per hectare (61.98 kg) was recorded in P_5_×P_1_. Positive standard heterosis was observed for key traits over check variety and mostly developed hybrids showed better performance in all traits compared to check variety. Pearson correlation and PCA analysis revealed key trait associations and genetic variation among sweet pepper hybrids, aiding breeding strategies. Considering morphological, physiological, yield potential and heterosis, P_5_×P_1_, P_7_×P_1_, P_6_×P_5_, P_3_×P_2_, and P_5_×P_3_ are recommended for further evaluation and breeding programs.

## Introduction

Sweet pepper (*Capsicum annuum* L.), belonging to the Solanaceae family, is a commercially valuable vegetable widely grown around the world, especially in temperate climates. It is the fourth most widely cultivated vegetable among the Solanaceous crops [1]. This diploid species possesses a chromosome number of 2n = 24 [2] and is believed to have originated in the tropical regions of South America [3]. It is now widely grown in Central and South America, including Costa Rica, Bolivia, Peru, and Mexico, as well as across most European countries, Hong Kong, and India [4]. It has a different color range from green to red, yellow, orange, purple and black which can be eaten as cooked or raw with different food items, as well as in salad. This crop has a consistently increasing international market and world production has expanded tremendously over the last few decades [5]. Sweet pepper is not only an economically significant vegetable crop but also offer high nutritional value. Sweet pepper is a rich source of vitamins, lycopene, xanthophylls, carotenoids, dietary fiber, and essential minerals [6]. According to different studies, 100 grams of the edible portion of sweet pepper contains approximately 8,493 IU of vitamin A, 283 mg of vitamin C, 13.4 mg of calcium, 14.9 mg of magnesium, 28.3 mg of phosphorus, and 263.7 mg of potassium [7, 8].

In Bangladesh, sweet pepper is commonly referred to as capsicum and is regarded as a minor vegetable crop [9]. While capsicum is a key summer crop in temperate regions, recent efforts have been made to cultivate it in Bangladesh. Presently, a few progressive farmers grow capsicum on a limited scale to fulfill the demand in major metropolitan areas [10]. In the year 2018-2019, about 11 MT of sweet pepper was produced from an area of 8 acres in Bangladesh [11]. This production is not sufficient to meet the ever-increasing demand in Bangladesh, and there is a huge potentiality to expand the area under cultivation of sweet pepper. Moreover, there is also exist an export market for this high value crop. Although sweet pepper production is steadily rising in Bangladesh, farmers frequently face challenges in selecting appropriate varieties. To date, the Bangladesh Agricultural Research Institute (BARI) has developed only two open-pollinated sweet pepper varieties suited to local growing conditions: BARI Mistimarich-1 and BARI Mistimarich-2 [12]. However, due to limited availability of these seeds, farmers largely depend on imported seed sources [13]. Most of the imported varieties are not adapted to Bangladesh conditions and the resulting yield is affected negatively since sweet pepper is highly sensitive to environmental conditions. [14]. Therefore, the successful production of this crop to meet the ever-increasing demand requires not only the development of new hybrids but also their performance and heritability study in terms of desired characteristics [15]. Golakia and Maken [16] reported that the improvement of yield and quality in any crop can be achieved by selecting genotypes with desirable traits naturally present or through genetic manipulation involving diverse parents through hybridization. Commercial hybrids have consistently demonstrated their superiority in terms of marketable fruit yield and fruit quality compared to open-pollinated cultivars, primarily due to the phenomenon of heterosis. Heterosis, often referred to as hybrid vigor, denotes the enhanced performance of F_1_ hybrids over their parent varieties in terms of yield and other desirable traits. Heterosis has been widely utilized in agriculture to enhance yield and adaptability in hybrid varieties across various crop species [17].

Agrawal [18] reported, when a hybrid surpasses the mid-parent value, which is essentially the average performance of its two parent plants, it is referred to as exhibiting heterosis. However, if the performance of the F_1_ hybrid exceeds that of the better parent, it is termed heterosis. It’s important to note that the practical utilization of heterosis in plant breeding is only viable when the vigor of the F_1_ hybrid significantly surpasses that of both the existing commercial hybrid and the superior parent plant. Although each parent selected for a specific crossing program has the potential to contribute desirable characteristics to its new generation, most breeders want to have a piece two of more elaborative information and scientific understanding of the heritability potential for the expression of these characteristics. This will ultimately allow the breeders to more effectively utilize parental lines to produce better hybrids expressing enhanced attributes promptly. Currently, the evaluation of different physiological and morphological characteristics, yield and various phytochemicals in sweet pepper is of most interest among the scientists [19]. Butcher et al. [15] showed that certain parent combinations were effective in producing F1 hybrids with improved fruit traits, such as weight, length, diameter, and higher phytochemical content. Identifying cross combinations with significant positive heterosis is important for boosting productivity and increasing farmers’ income. Developing new hybrids with better fruit traits, higher nutritional quality, and rich phytochemicals can help to create improved cultivars for human consumption. This study aimed to evaluate the performance of sweet pepper hybrids and check varieties based on their morphological, physiological, and yield-related traits. Additionally, the investigation aimed to quantify the standard heterosis of hybrids and check varieties, thereby providing insight into the degree of superiority or inferiority of hybrids relative to their reference varieties.

## Materials and Methods

### Experimental site

The study was carried out at the research field of the Department of Horticulture, Sylhet Agricultural University, from October 2022 to June 2023. The experimental site is situated in northeastern Bangladesh, within Agro-Ecological Zone (AEZ) 20, and lies between 23.57° to 25.13° N latitude and 90.56° to 92.21° E longitude. This subtropical region experiences distinct seasonal climate variations. The soil at the experimental site is highly acidic, with a silty clay loam texture and a pH range of 4.7-6.9 [20, 21].

### Plant material and growing condition

Eight newly developed sweet pepper hybrids and two check varieties (Table 1) were selected based on high yield potential, diverse horticultural traits, strong combining ability, and desirable heterosis. The field experiment followed a Randomized Complete Block Design (RCBD) with three replications, while laboratory work used a Completely Randomized Design (CRD). Each replication included five plants, with over 80% plant survival. Plots measured 2.5 × 1 m, with 50 cm spacing between and within rows.

**Table 1.** A list of the chosen hybrids and the check varieties used in this experiment.

| Developed hybrids | Check varieties |
| --- | --- |
| $P_3 \times P_1$ | CV<sub>1</sub>=Sweet Beauty, Lal Teer, (Local hybrid) |
| $P_4 \times P_3$ | CV<sub>2</sub>= AVPP 0504, AVRDC, Taiwan (Parent variety) |
| $P_6 \times P_5$ | |
| $P_5 \times P_1$ | |
| $P_7 \times P_1$ | |
| $P_5 \times P_3$ | |
| $P_3 \times P_2$ | |
| $P_6 \times P_3$ | |

Seeds were soaked and sown on October 28, 2022, in trays filled with equal parts sandy loam soil and decomposed cow dung. Seedlings were transplanted into polybags on November 14 and kept under partial shade for 25 days. On December 20, 25-day-old seedlings with 5-6 leaves were transplanted to the field. Fertilization followed BARI’s recommended dose [22] (Suppl. Table 1). A nylon net (20 mm mesh) was installed to protect the plots, and standard intercultural operations were performed. Peppers were harvested at full maturity based on size, firmness, and gloss. Data was collected on various aspects of the sweet peppers, and each genotype was characterized according to the descriptor codes provided by IPGRI, AVRDC, and CATIE [23] for Capsicum (*Capsicum spp*.). For data collection, five plants were randomly chosen from each treatment and replication. Information on morphological, physiological, and yield parameters was recorded.

### Determination of chlorophyll and carotenoids content of leaves

A 0.1-gram sample of fresh sweet pepper leaves is blended with 6 milliliters of 80% acetone. After blending, the mixture is centrifuged at 8,000 rpm for 25 minutes at 4°C. The resulting liquid is used to measure chlorophyll and carotenoid levels. Absorbance is read at 470, 645, and 663 nm using a spectrophotometer (Model-UV-1900i, Shimadzu, Japan). The amounts of chlorophyll a, chlorophyll b, and total chlorophyll are then calculated using a modified version of the Lichtenthaler and Wellburn formulas from [24].

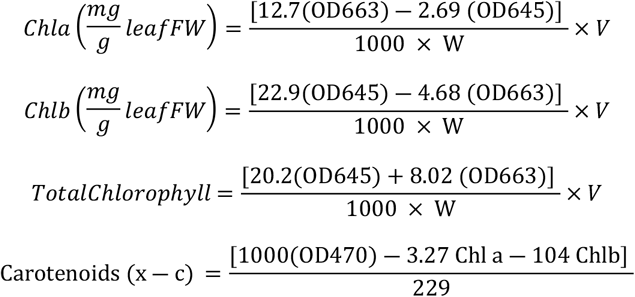

Where,

OD = Optical Density

V = Volume of sample

W = Weight of sample

### Estimation of Heterosis

Percent standard heterosis (S) for each character calculated by using this formula

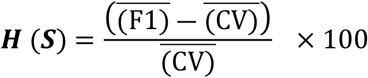

Where,

F_1_ = Mean value of each hybrid;

CV = Mean value of the check variety

The mean error variance from the combined analysis of variance for the check variety and F1 hybrids was used to calculate the standard error (SE) of the difference. Mean values across replications were compared. To estimate standard heterosis, the difference between F1 hybrids and the check variety was considered. If the difference exceeded the critical difference (CD), it was regarded as significant; otherwise, it was not.

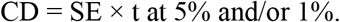

### Statistical Analysis

The collected data were analyzed using STATISTICA 10, R, and OriginPro software. Mean performances were compared through an F-test, and the coefficient of variation (CV %) was computed to assess variability.

## Results

### Morpho-physiological and yield characteristics of sweet pepper (F_1_ hybrids)

#### Classification of different morphological traits

A diverse range of morphological traits was observed in our research field (Table 2), including growth habit (prostrate, intermediate, and erect), branching pattern (sparse, intermediate, and dense), and stem color (green and light green). Variations were also noted in nodal anthocyanin pigmentation (green, purple, and light purple), leaf shape (ovate-lanceolate and deltoid), and flower position (intermediate and erect). Fruit morphology exhibited considerable diversity, with fruit shapes categorized as elongate, almost round, triangular, and blocky. Fruit color at the intermediate stage varied among yellow, green, purple, and light green, while mature fruit color ranged from lemon-yellow, light red, red, and dark red to light purple-orange. Additionally, fruit shape at the blossom end was classified as pointed, blunt, sunken, or a combination of sunken and pointed, whereas fruit shape at the pedicel attachment was identified as truncate, cordate, or lobate. The presence or absence of fruit blossom end appendages was also recorded from the hybrids and check varieties.

**Table 2.**
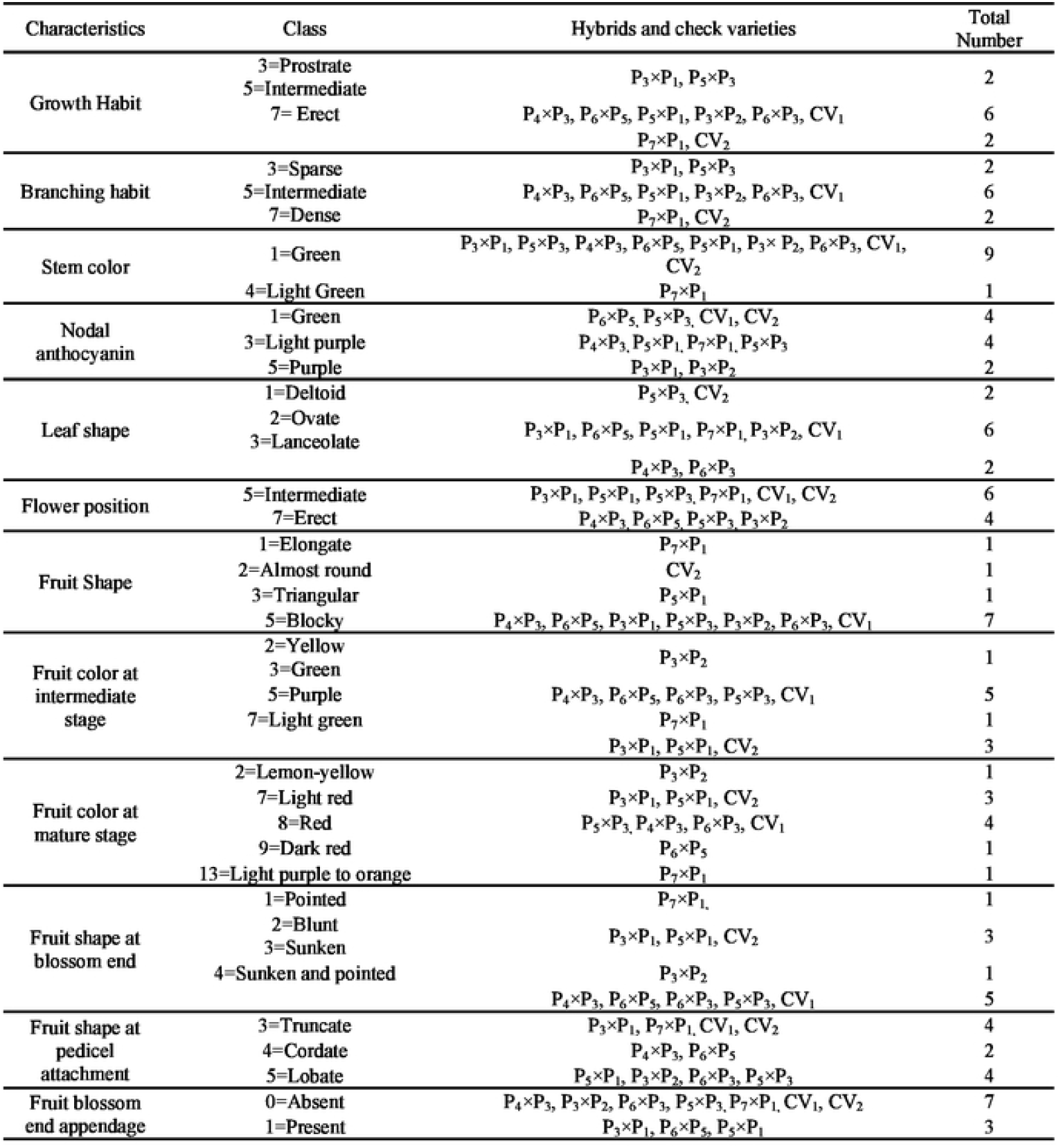
Classification and Morphological Characteristics of Sweet Pepper Hybrids and Check Varieties.

#### Morphological traits

Among the hybrids and check varieties, the tallest plants were recorded in CV_2_ (58 cm), P_3_×P_1_ (55.13 cm), and P_6_×P_5_ (55 cm). In contrast, the shortest plant (44 cm) was observed in CV_1_. The hybrid P_7_×P_1_ and P_3_×P_2_ exhibited the maximum stem diameter at 0.95 cm and 0.91 cm consecutively. In contrast, the minimum stem diameter was observed in CV_2_ and CV_1_ at 0.43 cm and 0.53 cm consecutively. The experimental results revealed that the developed variety P_7_×P_1_ had the highest number of branches per plant (15.13), while the lowest number was observed from CV_1_ (5), CV_2_ (5), P_3_×P_1_ (5.88) and P_5_×P_1_ (5.50). The highest number of leaves per plant, 64.63, 63.38, 61.75, and 61.75, was observed in P_6_×P_5_, P_3_×P_1_, P_4_×P_3_, and P_7_×P_1_, respectively. The lowest number of leaves per plant was recorded in CV_1_ (33.25). The highest plant surface area was observed in P_7_×P_1_ (1916 cm^2^) and CV_1_ (1743.5 cm^2^), while the lowest plant surface areas were found in P_6_×P_3_ (982.1 cm^2^) and P_3_×P_2_ (1081.8 cm^2^) (Suppl. Table 2).

#### Physiological traits

Most of the hybrids exhibited SPAD values similar to the check variety CV_2_, which recorded the highest value in the current study. The observed variation was attributed to genetic differences and other factors. The highest leaf chlorophyll content was observed in the parental variety CV_2_ (1.55 mg/g fresh leaf), whereas the check variety CV_1_ exhibited the lowest value (0.33 mg/g fresh leaf). The hybrids displayed intermediate chlorophyll content between CV_1_ and CV_2_. Similarly, the highest carotenoid concentration was recorded in the check variety CV_2_ (7.16 mg/g), while the lowest concentration was found in the check variety CV_1_ (1.55 mg/g). The carotenoid content of all hybrids ranged between those of CV_1_ and CV_2_ (Suppl. Table 2). Notably, most hybrids exhibited chlorophyll values comparable to CV_2_, which recorded the highest chlorophyll content.

#### Flowering traits

The hybrids P_3_×P_1_, P_4_×P_3_, P_5_×P_3_, and the check variety CV_2_ exhibited early flowering at 30.33, 32.64, 33.62, and 33.35 days, respectively, while the remaining hybrids showed delayed flowering. Similarly, early fruit setting was observed in the hybrids P_3_×P_1_ (38.33 days), P_4_×P_3_ (41.35 days), P_5_×P_3_ (41.62 days), and P_3_×P_2_ (40.64 days), whereas the other hybrids exhibited delayed fruit setting (Suppl. Table 3). Notably, early fruit setting was more pronounced in the hybrids compared to the check variety.

**Table 3.**
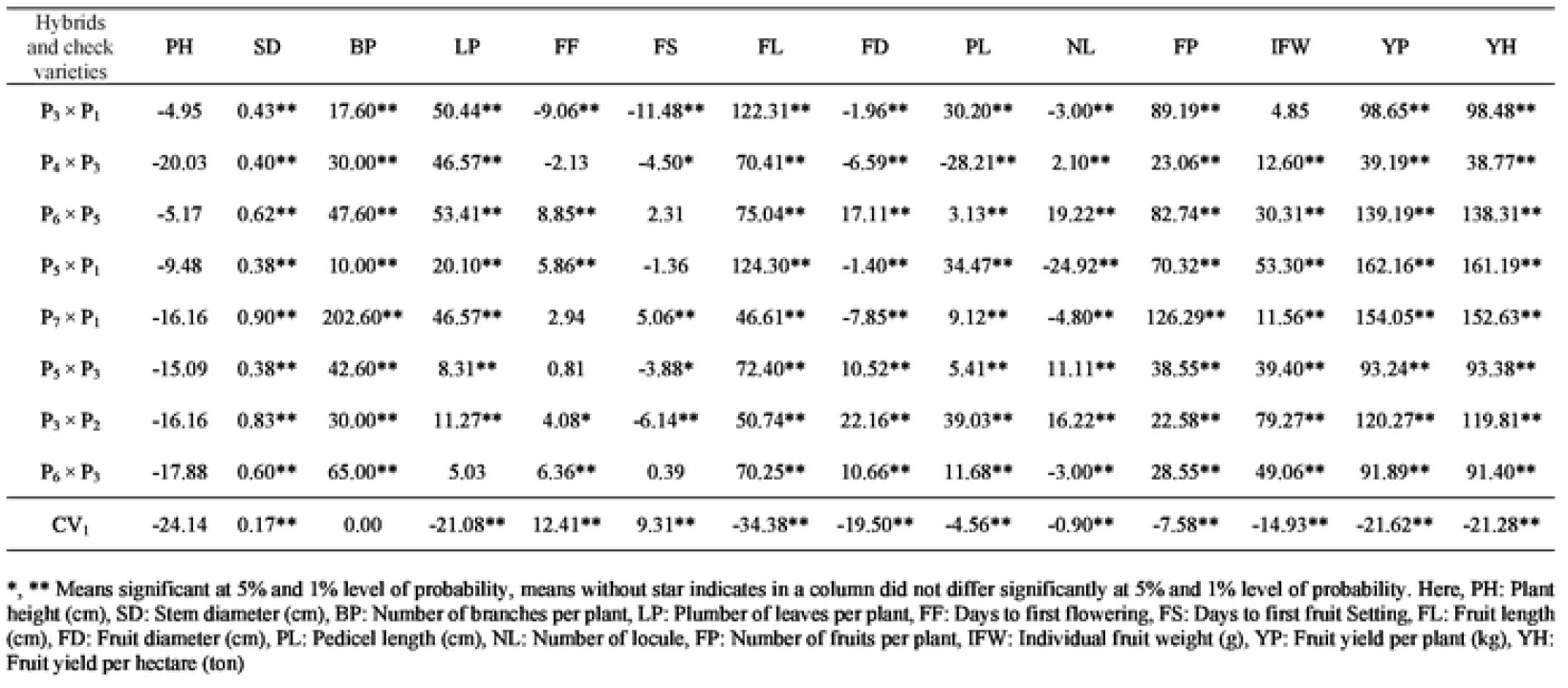
Percent or standard heterosis of hybrids over check variety (CV2) for morphological and yield characteristics of sweet Pepper.

#### Yield attributes

The fruit length of sweet pepper ranged from 3.97 to 13.57 cm. Among the different hybrids and check varieties, the hybrids P_5_×P_1_ and P_3_×P_1_ had the longest fruit length at 13.57 cm and 13.45 cm. Conversely, the shortest fruit length at 3.97 cm was observed in check variety CV_1_. The hybrids P_3_×P_2_ displayed the largest fruit diameter measuring 8.71 cm, while the smallest fruit diameter of 6.74 cm was observed in the check variety CV_1_. The hybrid exhibited larger mean fruit diameters compared to the check varieties. Among the hybrids and check varieties, the hybrids P_3_×P_2_ and P_5_×P_1_ exhibited the longest pedicel length for fruit measuring 4.88 cm and 4.72 cm, whereas the check variety CV_1_ and CV_2_ had the shortest pedicel length at 3.35 cm and 3.51cm respectively. From the hybrids and check varieties, the check variety CV_2_ had highest recorded pericarp thickness value (1.36 cm), whereas the P_6_×P_3_ cross variety exhibited the lowest pericarp thickness (0.633 cm) in the current study. In the study the P_6_×P_5_ and P_3_×P_2_ hybrids showed the maximum number of locule at 3.97 and 3.87 respectively, while the P_5_×P_1_ hybrids displayed the smaller number of locule at 2.50 (Suppl. Table 3). The results of our study revealed that the sweet pepper hybrid P_3_×P_2_ produced the highest number of seeds per fruit (183.30), whereas the check variety CV_1_ had the lowest seed count per fruit (52.33). The seed count for the hybrids P_5_×P_3_, P_5_×P_1_, P_3_×P_1_, and P_6_×P_5_ was recorded as 176.67, 158.48, 143.86, and 134.80, respectively (*Suppl. Figure 1A*). Regarding seed weight, the hybrid P_6_×P_5_ exhibited the highest seed weight (9.30 g), while the lowest seed weight was observed in the hybrid P_6_×P_3_ (7.30 g) (*Suppl. Figure 1B*). Notably, the hybrids demonstrated a significantly higher seed count per fruit compared to the check varieties. This finding has important implications for future cultivation, as increased seed production enhances propagation potential for subsequent generations.

The sweet pepper hybrids P_7_ × P_1_ displayed the utmost number of fruits per plant 14.03, whereas check variety CV_1_ and CV_2_ exhibited the lowest 5.73 and 6.20 number of fruits per plant respectively (*Figure 1*). The highest sweet pepper individual fruit weight (214.51 g) was recorded from hybrids P_3_×P_2_, while check variety CV_1_ exhibited the lowest individual fruit weight 101.80 g (*Figure 2*). The highest sweet pepper fruit yield per plant (1.94 kg) and (1.88 kg) was observed from the hybrid P_5_×P_1_ and P_7_×P_1_ respectively, whereas the lowest fruit yield per plant (0.58 kg) was recorded from the check variety CV_1_ (*Figure 3*). The sweet pepper hybrids P_5_×P_1_ and P_7_×P_1_ demonstrated the highest fruit yield per plot (15.49 kg) and (14.99 kg) respectively, while the lowest fruit yield per plot (4.67 kg) was recorded from the check variety CV_1_ (Suppl. Figure 2A). The highest sweet pepper fruit yield per hectare (61.98 ton) and (59.95 ton) was observed from hybrids P_5_×P_1_ and P_7_×P_1_ respectively, whereas the lowest fruit yield per hectare (18.68 ton) was recorded from check variety CV_1_ (Suppl. Figure 2B).

**Figure 1.**
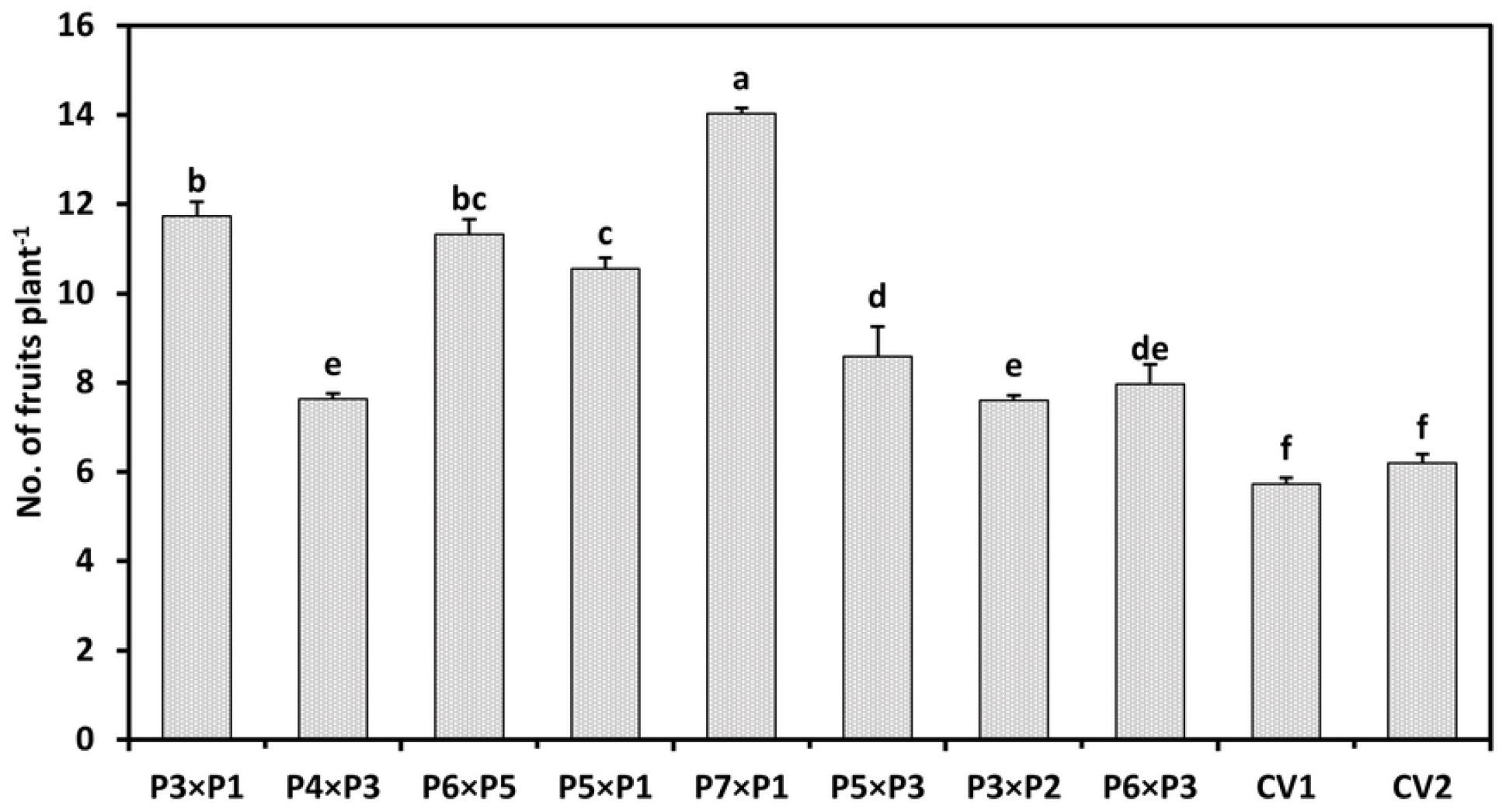

**Figure 2.**
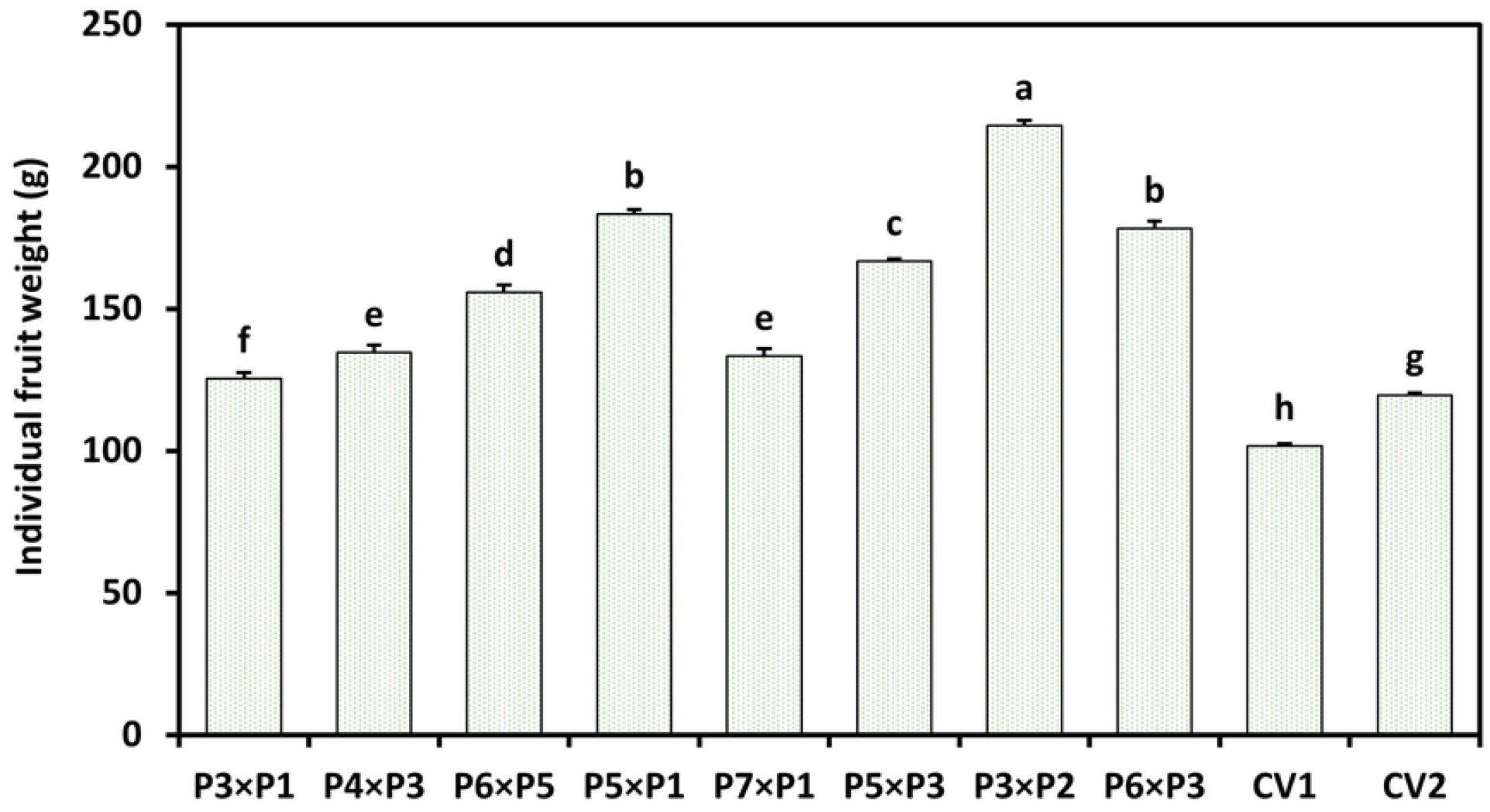

**Figure 3.**
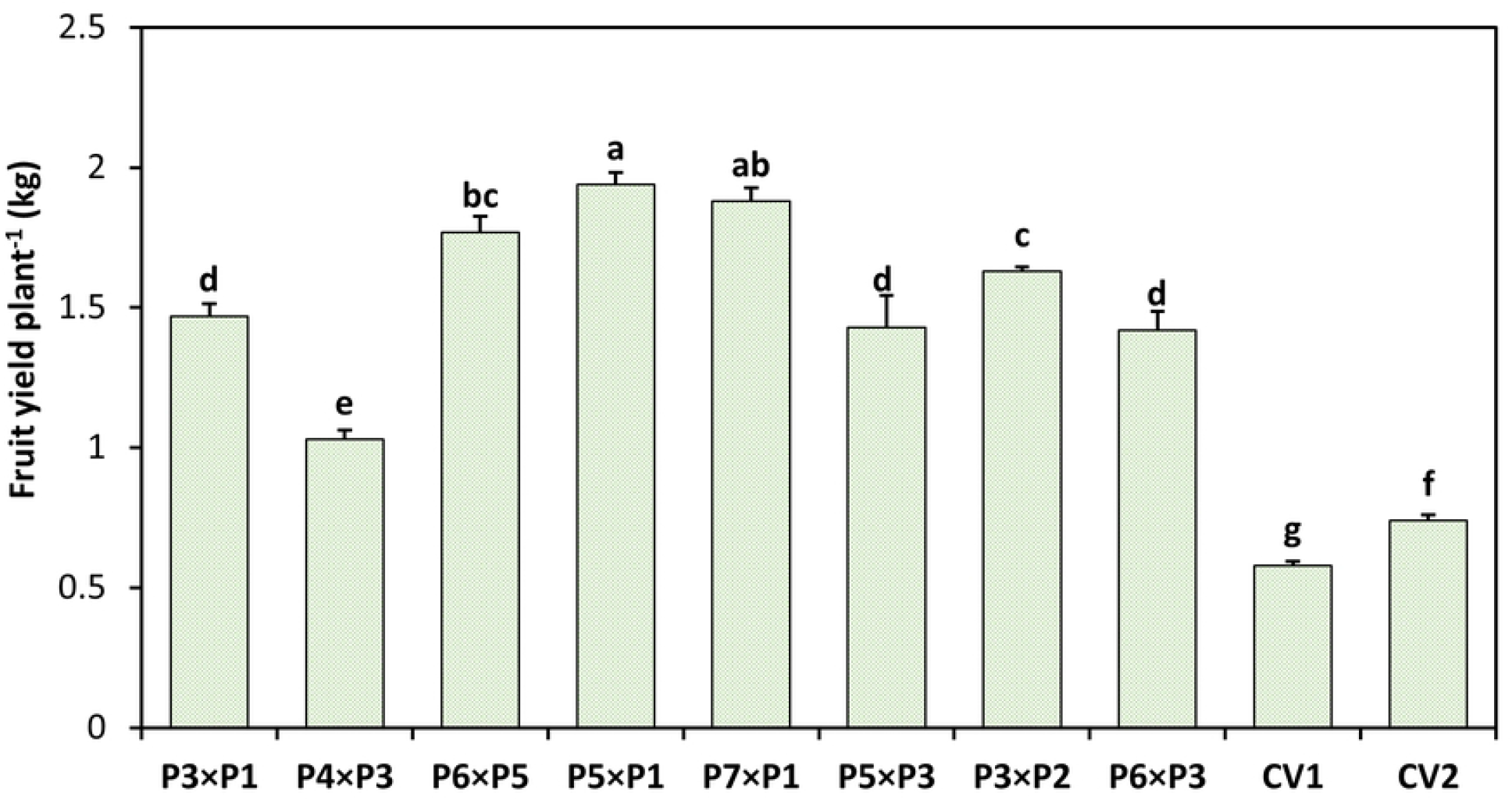

### Trait Association

The Pearson correlation analysis revealed a total of 231 trait associations, among which 39 were statistically significant (*p*< 0.05) (*Figure 4*). Among these, 38 showed a positive correlation, while only one displayed a significant negative correlation. Strong positive correlations were observed between key yield-related traits such as fruit yield per plant (FYP), fruit yield per plot (FYPl) (r = 1.00, *p*< 0.001) and number of fruits (NF) with fruit yield per ton (FYT) (r = 0.80, *p*< 0.001). Morphological traits such as number of branches (NB) exhibited a positive correlation with stem diameter (SD) (r = 0.71, *p*< 0.05). Pericarp thickness (PT), chlorophyll SPAD value (CS), leaf chlorophyll (LC) and carotenoids (LCa) were positively correlated with plant height (PH). The only significant negative correlation was observed in fruit length (FL) with days to fruit setting (DFS) (r= −68, p < 05). These findings highlight key trait associations that can be utilized for breeding strategies aimed at enhancing sweet pepper yield and quality.

**Figure 4.**
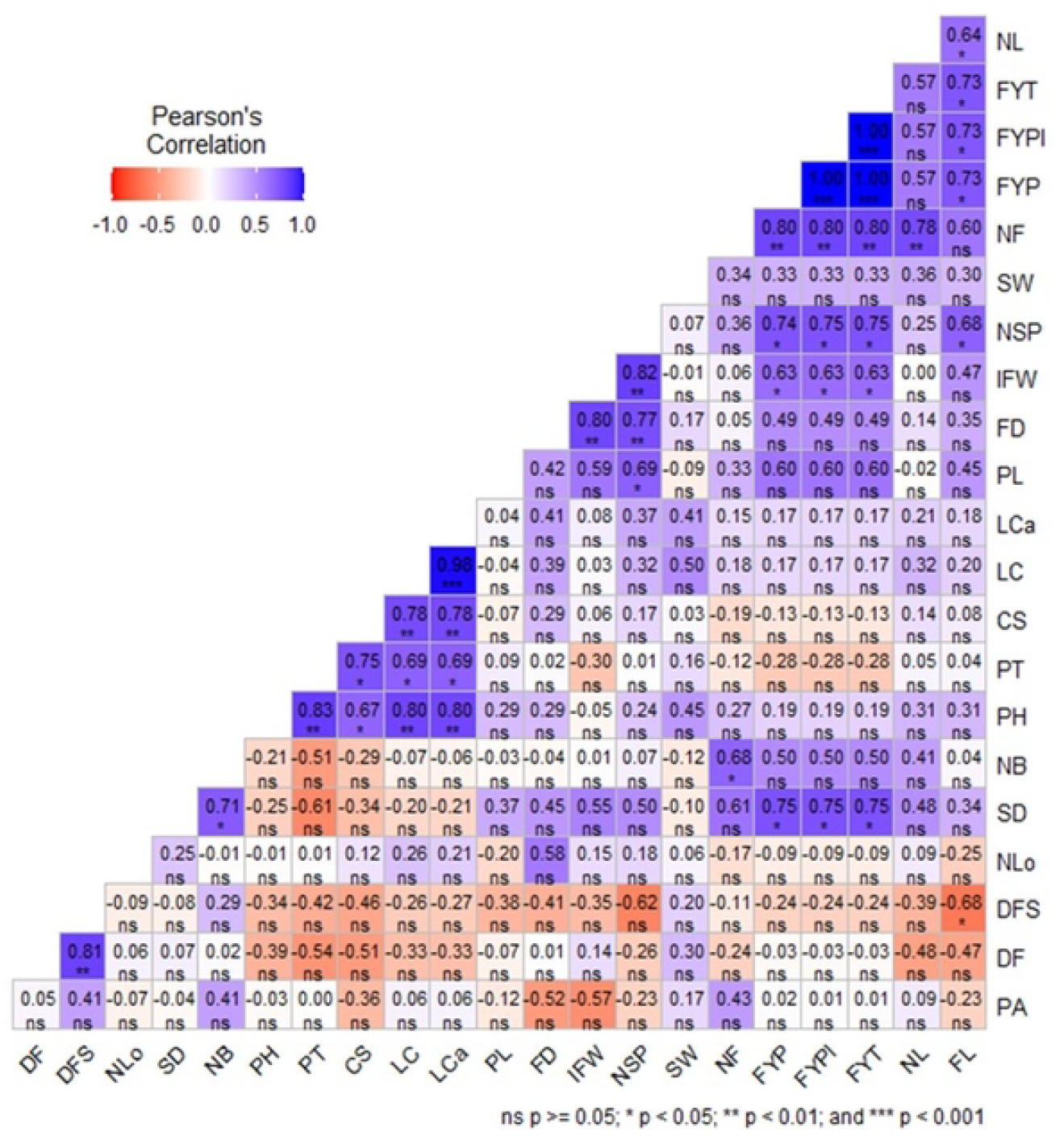

The PCA biplot (*Figure 5*) illustrates the relationships among morphological, physiological, and yield traits of sweet pepper hybrids and check varieties. The first two principal components (PC1 and PC2) explain 39.38% and 27.82% of the total variance, respectively, accounting for a cumulative 67.2% of the total variability. The trait vectors indicate the contribution and grouping of different characteristics. Yield-related traits such as fruit yield per plant (FYP), fruit yield per ton (FYT), and number of fruits (NF) are clustered closely, indicating a strong association. Morphological traits like plant height (PH), stem diameter (SD), and number of branches (NB) are also positioned in proximity, suggesting similar influences. The distribution of hybrids (red dots) suggests genetic variation in performance across different genotypes. Hybrids such as P_3_×P_2_ and P_5_×P_3_ are positioned far from the origin, indicating distinct trait expressions compared to other combinations. The placement of hybrids relative to trait vectors provides insights into their phenotypic performance, useful for breeding and selection programs.

**Figure 5.**
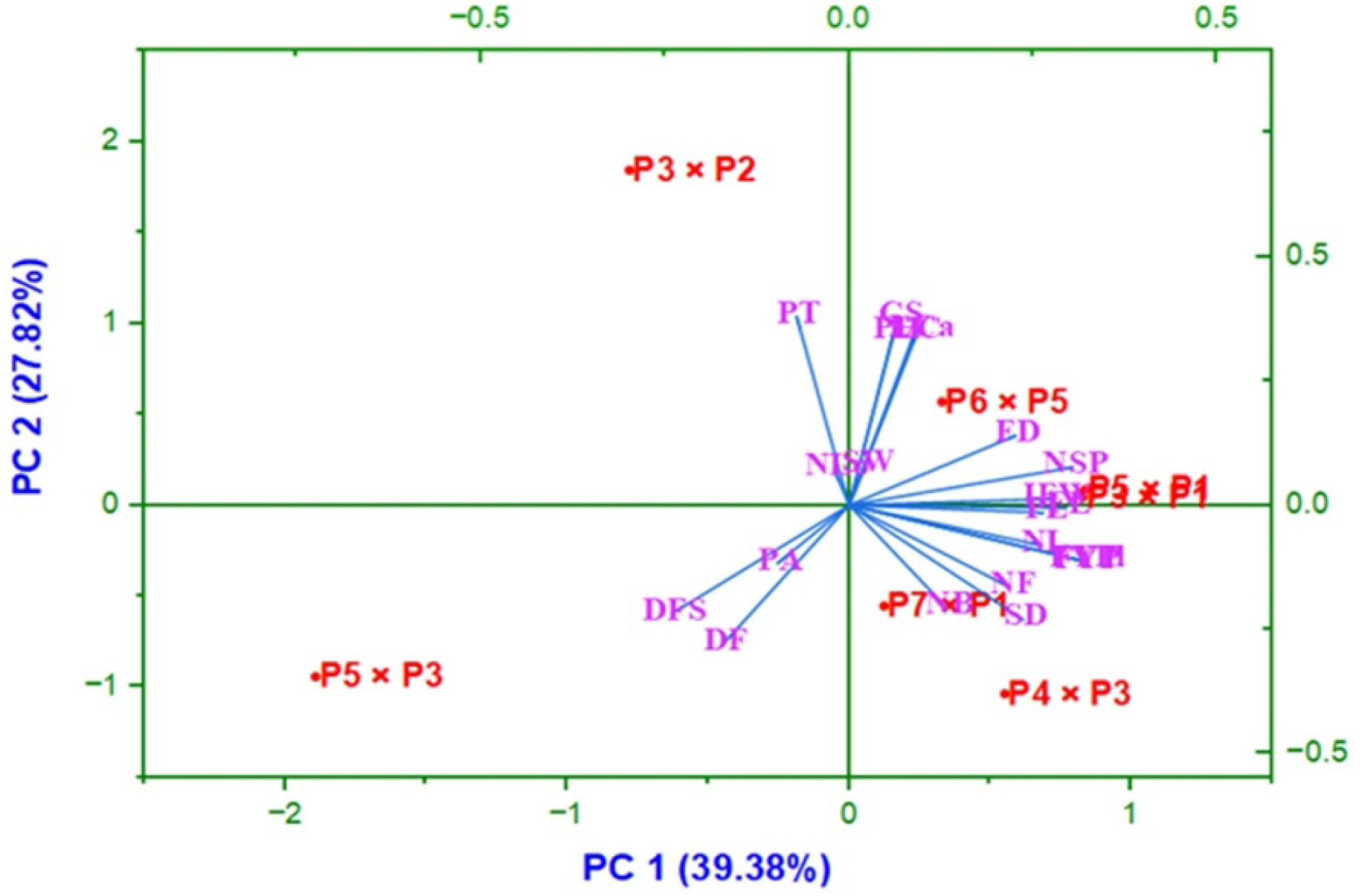

### Analysis of Variance (ANOVA)

Sweet pepper hybrid and check types were compared using analysis of variance (ANOVA), which revealed notable differences for several characteristics (Suppl. Table 4). The treatment effects were highly significant (p < 0.01) for plant height, stem diameter, number of branches and leaves per plant, fruit length, fruit diameter, pedicel length, individual fruit weight, and yield per plant and hectare. Days to first flowering and fruit setting showed moderate significance (*p*< 0.05), while replication effects were mostly non-significant. The high significance in yield-related traits indicates considerable genetic variation among the hybrids, suggesting the potential for selecting superior varieties for commercial cultivation.

### Estimation of Standard Heterosis

The standard heterosis percentage for various morphological and yield traits in sweet pepper hybrids exhibited significant variations with positive and negative standard heterosis. The hybrid P_5_×P_1_ demonstrated the highest positive standard heterosis for fruit yield per plant (162.16%) and per hectare (161.19%), followed by P_7_×P_1_ with 154.05% and 152.63%, respectively. The number of branches per plant showed the maximum positive heterosis in P_7_×P_1_ (202.60%), while P_5_×P_1_ had the lowest significant increase (10.00%). P_3_×P_1_ exhibited the greatest standard heterosis in number of leaves (50.44%) and fruit length (122.31%), whereas P_3_×P_2_ showed the highest heterosis for fruit diameter (22.16%) and pedicel length (39.03%). P_6_×P_5_ exhibited superior standard heterosis for individual fruit weight (30.31%) and number of locules (19.22%). First flowering and fruit setting also showed positive standard heterosis in developed hybrids. Some negative standard heterosis recorded from developed hybrids. It indicated that some trait showed good result over CV_2_. The local check variety CV_1_ consistently showed negative heterosis across most traits, indicating the superior performance of newly developed hybrids. These findings suggest that P_5_×P_1_, P_7_×P_1_, P_6_×P_5_, and P_3_×P_2_ can be considered promising candidates for commercial cultivation based on their significant heterotic advantages (Table 3).

## Discussion

Sweet pepper exhibits considerable genetic variability across morphological, physiological, and yield-related traits, which plays a pivotal role in breeding programs aimed at developing improved cultivars. The present study evaluated a diverse set of sweet pepper hybrids and check varieties, revealing significant differences in key agronomic traits such as plant height, branching pattern, flowering behavior, fruit characteristics and yield attributes. These findings are consistent with previous studies and provide valuable insights into the genetic potential of sweet pepper hybrids for enhancing productivity and quality. Sweet peppers predominantly exhibit an intermediate growth habit and varying intensities of nodal anthocyanin, ranging from dark to light purple [25]. Most genotypes showed an intermediate branching habit, with only one displaying a sparse habit and two exhibiting dense branching. The majority had green fruit color at the intermediate stage, while all except CKN-8 and CKN-11 lacked a fruit blossom end appendage [26]. A dominant green stem color was recorded in 66.66% of the germplasm, with 28.57% showing green with purple streaks and 4.76% light green with purple streaks. Additionally, 52.38% had an intermediate flower position, while 38.0% and 9.52% exhibited pendant and erect pedicel positions, respectively [27]. Capsicum spp. displays a range of fruit shapes, including campanulate, blocky, ovoid, round, almost round, linear, triangular, rectangular, and elongate, with mature colors such as red, orange, yellow, brown (chocolate), and cream [28]. A study on 43 sweet pepper progenies revealed variations in fruit shapes, with 25 showing a sunken blossom end, 8 a pointed end, and 13 a blunt end. Progeny also exhibited different pedicel attachment shapes: 10 truncate, 20 cordate, and 12 lobate [29].

At the 50% flowering stage, Ahmed et al. [30] reported that the tallest plant height of sweet pepper, 52.44 cm, was achieved with 28-day-old transplants (T2), while the shortest height, 39.66 cm, was observed with 48-day-old transplants (T6). Similarly, Tuckeldoe et al. [31] conducted a study on sweet pepper, where the stem diameter ranged from 0.82 to 1.18 cm in the 2021 season and from 0.89 to 1.02 cm in the 2022 season. Maniutiu et al. [32] reported that the sweet pepper plant exhibits diverse branching habits, and it is suggested that controlling fruit development can be achieved by maintaining a branching pattern ranging from 1 to 4 branches per plant, which can contribute to an extended fruiting period. In a related study, Al-Nassrallah and Al-Asadi [33] reported the highest number of primary branches per sweet pepper plant at 11.24, with the lowest recorded at 7.32 branches per plant. Roy et al. [34] reported a wide range in the number of leaves per sweet pepper plant, with values varying from 43.67 in 8/4 × Royal Wonder to 82.75 in Baby Bell × C/4. Barthlott et al. [35] reported that dissimilarity in plant surface area plays a crucial role in environmental interactions, particularly for sessile organisms with extensive functional surfaces, such as plants. In a previous experiment, Messi showed the highest relative chlorophyll content at 50.08, followed by Maria (44.8) and BARI Mistimorich-1 (41.5). In contrast, the lowest relative chlorophyll content was observed in Thunder (29.33), followed by Green Bell (34.5) and Dream (37.4), according to the findings of Ghosh et al. [36]. Dhaliwal et al. [37] reported that in Capsicum spp., total chlorophyll content in leaves ranged from 1.17 to 1.67 mg/g fresh weight.

Furthermore, the majority of the developed hybrids demonstrated carotenoid concentrations similar to those of the check variety with the highest recorded value. These findings align with those of Hegazi et al. [38], who reported comparable carotenoid concentrations in their study. The combined mean response indicates that the Jeju and Markofana varieties of sweet pepper achieved 50% flowering earliest at 40 days, while Jeju exhibited relatively delayed flowering at 43 days, a trait beneficial for early fruit harvest. Additionally, Odaharo reached 50% fruit setting the earliest at 67 days, whereas Jeju attained this stage later at 76 days, as reported by Chernet and Zibelo [39]. Galal et al. [40] reported that cross genotypes of sweet pepper exhibited a fruit length of 7.93 cm (P1×P2), while the lowest fruit length was 5.17 cm (P2×P5) and the cross genotype of sweet pepper P2×P3 had a fruit diameter of 6.10 cm, while the lowest fruit diameter of 3.90 cm was observed in P1×P5. Similarly, Sharma et al. [41] reported that the highest pedicel length of sweet pepper was recorded in the PBC-631 × YW hybrid genotype (3.258 cm), while the lowest pedicel length was observed in the IHR-546 × CW hybrid genotype (1.765 cm) and the sweet pepper genotype PBC-631 × YW hybrid exhibited the maximum pericarp thickness (0.501 cm), while the PBC-631 × CW hybrid showed the minimum pericarp thickness (0.285 cm). Danojevic and Medic-Pap [42] conducted a study where they noted that the sweet pepper hybrid genotypes exhibited a range of locule numbers, with the highest being 3.33 and the lowest recorded at 2.35. A comparable study by Shrestha et al. [43] reported that while SP6, SP35, and SP39 genotypes of sweet pepper produced seedless fruits, SP43 yielded the highest seed count per fruit, averaging 147, highlighting the significance of seed production for crop sustainability. Furthermore, variations in thousand-seed weight were observed in the present study. A similar trend in sweet pepper was reported by Bhattarai et al. [44], where the cultivar ‘California Wonder’ had the highest 1000-seed weight (7.44 g), while HRDCAP-001 recorded the lowest (6.00 g). In their study, Sharma et al. [45] documented the highest count of sweet peppers in KandaghatSel-1 at 9.30 and the lowest in Solan Wonder at 3.83, as well as the highest individual fruit weight in PRC-1 at 80.82 g and the lowest in Solan Pepper at 37 g, highlighting significant distinctions among genotypes. Similarly, Galal et al. [40] reported the highest fruit yield per plant (1.65 kg) from P4×P5 and the lowest (0.77 kg) from P3. Furthermore, the highest fruit yield per plant (693.10 g) in PRC-1 and the lowest (163.96 g) in Solan Wonder from their research. These collective findings underscore the diversity in fruit yield among different varieties. Consistent with these findings, both Galal et al. [40] and Sharma et al. [41] observed similarities in their research results, further supporting the notion of significant variation in fruit yield among different sweet pepper genotypes. Similarly, findings from Bhattarai et al. [44] align with these results, indicating the highest yield per hectare from the HRDCAP-001 genotype (68.3 t/ha) and the lowest from the HRDCAP-004 genotype (40.3 t/ha).

Similar trends have been reported in previous studies, where heterosis magnitudes for plant height, branch number, fruit length, and pedicel length varied considerably across hybrids [1, 46, 47]. Consistent with Roy et al. [34], positive heterosis for early flowering traits was observed, highlighting the potential for improved earliness in breeding programs. While some hybrids exhibited negative heterosis in specific traits, these variations align with prior findings on heterotic effects in *Capsicum annuum* L. [48, 49]. Furthermore, fruit number per plant and fruit weight showed considerable heterotic advantages, supporting their relevance in hybrid selection [50, 51]. Overall, the study confirms the heterotic potential of selected hybrids, reinforcing their suitability for large-scale cultivation and genetic improvement in sweet pepper breeding.

## Conclusion

This study evaluated the morphological, physiological, and yield characteristics of sweet pepper hybrids, highlighting significant genetic variations. The hybrids P3×P1 and P3×P2 exhibited early flowering and fruit setting. Among the hybrids, P_7_×P_1_ had the highest anthocyanin content, while P_3_×P_2_ had the highest Vitamin C content. The hybrids P_5_×P_1_, P_7_×P_1_, P_6_×P_5_, P_3_×P_2_, and P_5_×P_3_ performed better than the check variety in terms of morphological, physiological, yield characteristics. The heterosis study also confirmed that mostly hybrids showed better result compared to the check varieties for morphological, physiological and yield-related traits. The newly developed hybrids suggested to maintaining properly for further use in breeding program and it should be taken under considerations for multi locations trial.

